# MassNet: billion-scale AI-friendly mass spectral corpus enables robust *de novo* peptide sequencing

**DOI:** 10.1101/2025.06.20.660691

**Authors:** A Jun, Xiang Zhang, Xiaofan Zhang, Jiaqi Wei, Te Zhang, Yamin Deng, Pu Liu, Zongxiang Nie, Yi Chen, Nanqing Dong, Zhiqiang Gao, Siqi Sun, Tiannan Guo

## Abstract

Breakthroughs in artificial intelligence (AI) for natural language processing and computer vision have been largely driven by high-quality, large-scale datasets such as OpenWebText and ImageNet. Inspired by this, we present MassNet, a foundational resource for proteomics designed to accelerate deep learning applications. MassNet is the largest known corpus of data-dependent acquisition (DDA) mass spectrometry (MS) data, derived from ~30 TB of raw files and comprising 1.54 billion MS/MS spectra, resulting in 558 million peptide-spectrum matches (PSMs) across 35 species, including animals, plants, and microbes. Within the human subset, MassNet includes more than 1.7 million precursors and 19,966 proteins, covering 98% of annotated human proteins. To enable efficient AI training, we developed the Mass Spectrometry Data Tensor (MSDT), a structured format based on Parquet that enables standardized, high-performance batch access and seamless integration with GPU and TPU platforms for distributed training. We further extended MassNet to support *de novo* peptide sequencing, which infers peptide sequences directly from MS/MS spectra without reference databases, and is critical for discovering novel proteins, characterizing non-model organisms, and identifying post-translational modifications (PTMs). We introduce XuanjiNovo, a non-autoregressive Transformer model that leverages a curriculum learning strategy to enhance training stability. By dynamically adjusting learning difficulty based on model performance, XuanjiNovo achieves smooth convergence on complex, multi-distributional data without manual hyperparameter tuning. Trained on 100 million PSMs from the MassNet, it consistently outperforms state-of-the-art methods across diverse benchmarking tasks. Peptide recall exceeds 0.8 on the *Bacteroides thetaiotaomicron* and *Zea mays* datasets. On human data acquired using the Orbitrap Astral platform, XuanjiNovo achieves achieves 38.8% to 144.3% improvement over existing models. MassNet represents the first large-scale, standardized foundational dataset in proteomics, marking a critical milestone in the integration of artificial intelligence into proteomics research.

## 1 Introduction

Mass spectrometry (MS)-based proteomics has emerged as a pivotal technology in biomedical and life sciences, enabling the identification, quantification, and functional characterization of proteins in complex biological systems. Central to this field is data-dependent acquisition (DDA), the traditional and widely utilized method for MS data acquisition. In the past decade, proteomics data acquisition capabilities have advanced rapidly, leading to the continuous accumulation of large-scale datasets. For example, the PRIDE database, one of the world’s largest public repositories of MS data, currently hosts more than 42,000 datasets, encompassing a wide range of biological species, tissue types, disease states, and experimental designs [1]. Unlocking the full potential of these data for biological discovery increasingly depends on advanced computational tools, particularly those driven by artificial intelligence (AI).

AI-based models are progressively transforming MS data analysis. Unlike traditional rule-based or heuristic approaches, deep learning models can automatically learn complex, high-dimensional relationship from large-scale data [2]. This includes the recognition of subtle fragmentation patterns associated with post-translational modifications (PTMs) or non-tryptic cleavage events, without requiring manually predefined assumptions. Such data-driven capabilities allow deep models to deliver greater accuracy and flexibility in peptide identification and spectrum interpretation. Recent advances in deep learning have demonstrated remarkable breakthroughs across a range of proteomics tasks such as *de novo* peptide sequencing [3–7], peptide-spectrum matches (PSMs) rescoring [8–12], and various peptide property prediction tasks such as MS/MS spectrum prediction [13, 14], retention time (RT) estimation [13, 15, 16], and collision cross section (CCS) prediction [17]. Yet, as demonstrated in adjacent fields like natural language processing (NLP) and computer vision (CV), the power of AI models is inextricably linked to the quality, diversity, and structure of their training data [2]. In fields like NLP and CV, the emergence of large, well-annotated datasets, such as OpenWebText [18], WordNet [19], and ImageNet [20], have been instrumental in enabling self-supervised and supervised deep learning approaches to thrive. These datasets are not only extensive in scale but also meticulously curated, ensuring consistent data structures, rich annotations, and rigorous control of error rates. Proteomics, by contrast, lacks an equivalent foundation. While existing repositories have accumulated vast amounts of spectral data, they are not yet well-suited for artificial intelligence applications [21]. Specifically, they lack the structural consistency and completeness required for modern deep learning workflows. Key challenges include inconsistent data formats, variable data quality, incomplete metadata, and limited representation across experimental conditions and diverse species.

Some early community-scale initiatives, such as the Global Proteome Machine Database (GPMDB) [22], PeptideAtlas [23], the National Institute of Standards and Technology (NIST) [24], ProteomeTools [25], and jPOSTdb [26], played a pioneering role in the development of large-scale spectral libraries by systematically aggregating and curating high-quality MS/MS spectra for peptide identification and downstream applications. However, these spectral libraries often suffer from incomplete documentation of data sources and lack detailed traceability that links spectral library entries to the original raw data, database search results, and the software tools used in their construction. The emergence of MassIVE-KB [27] has largely addressed these issues by introducing a standardized and traceable data management mechanism, making it one of the most high-quality and representative human spectral libraries to date. MassIVE-KB continues to be updated and optimized, reflecting the ongoing demand for rigorous, large-scale human proteomic resources. However, most mainstream spectral libraries, including MassIVE-KB, remain heavily focused on human samples, with relatively limited attention given to proteomics data from other key model organisms. Yet species such as *Mus musculus, Drosophila melanogaster, Caenorhabditis elegans*, and *Saccharomyces cerevisiae*, play indispensable roles in a wide range of research areas, including biomedical science, developmental biology, drug discovery, and neuroscience. Their proteomes provide invaluable insights into evolutionarily conserved mechanisms, functional protein studies, and disease modeling. The lack of high-quality, species-diverse spectral resources has thus become a key bottleneck for broadening AI applications in proteomics beyond human data.

In addition to limited organismal coverage, existing spectral libraries also exhibit significant limitations in data representation. In most cases, peak lists are stored separately from their associated structural and identification information, such as peptide sequences, charge states, and modification sites. For example, spectral data are typically saved in formats like MGF or mzML, while search results are commonly stored in various unstructured or semi-structured formats, such as .msp [24], .sptxt [23, 27], or .tsv [9, 13]. These files often contain redundant and inconsistently annotated information, lacking a unified data schema or semantic standard. Such loosely organized formats significantly increase the complexity of parsing, integration, and downstream analysis, making them poorly suited for scalable computational workflows. Moreover, the file sizes typically range from hundreds of megabytes to several gigabytes, resulting in frequent I/O operations and slow data loading speeds. These limitations severely hinder large-scale parallel processing and significantly reduce the efficiency of deep learning model training on MS data. In contrast, tensor-based spectrum storage formats provide a powerful alternative. By encoding spectral information into compact two-dimensional tensors (e.g., m/z × intensity matrices), these formats support efficient batch loading, parallel processing, and GPU acceleration [28, 29]. They align naturally with deep learning model input requirements, significantly enhancing performance and scalability. Furthermore, tensorized data dramatically reduces memory footprint and I/O latency, while enabling seamless integration with modern AI frameworks. In essence, tensor-based libraries remove a major technical barrier, lowering both the computational and developmental costs of AI-powered proteomics.

Here, we introduce MassNet, a large-scale, AI-friendly dataset designed for deep learning applications in proteomics. It comprises 558 million PSMs, totaling 3.2 TB in size, and is derived from ~ 30 TB of raw DDA-MS data spanning 35 species and over 10 high-resolution MS platforms. Each spectrum includes detailed annotations such as project identifiers, instrument parameters, and search engine scores, ensuring full traceability and reproducibility for large-scale modeling tasks. All spectral and identification data are stored in a compact tensor format, capturing critical attributes such as m/z, intensity, and retention time in a two-dimensional representation suitable for deep learning. This design enables efficient model training, inference, and deployment in high-throughput environments. Training deep learning models on such large-scale datasets typically involves extensive hyperparameter tuning and iterative adjustments to training protocols, often spanning weeks to months [30]. This makes it challenging to directly apply existing models to complex, heterogeneous datasets like MassNet. To address this, we propose a robust and principled learning paradigm that achieves consistently successful optimization across diverse data distributions. Our approach begins with a simplified learning objective: the model is initially tasked with predicting only partial peptide sequences, with the remaining label tokens revealed as context. This curriculum learning [31] strategy significantly reduces early-stage learning difficulty and prevents training collapse, even in the presence of complex or imbalanced data. As training progresses, we continuously evaluate model performance on the given distribution and gradually increase task difficulty, enabling smooth and effective convergence. Building on the success of non-autoregressive *de novo* models such as PrimeNovo [7], we adopt a non-autoregressive Transformer [32] backbone that enables bi-directional context modeling in peptide sequences and significantly accelerates inference. By integrating additional post-processing modules such as sequence level self-refinement and PMC mass control unit, we develop a new model, XuanjiNovo. Trained on 100 million spectra from the MassNet dataset, XuanjiNovo consistently outperforms all existing *de novo* sequencing methods across multiple evaluation tasks, establishing a new state of the art in this field.

## 2 Results

### 2.1 Overview of the MassNet

The MassNet dataset comprises 27,643 DDA-MS files, primarily sourced from the PRIDE [1] and iProx [33] databases, encompassing ~ 30 TB of raw data and more than 1.5 billion MS/MS spectra (**Figure 1A**). Database searches were performed using two analysis tools: FragPipe [9, 34] and Sage [35], resulting in the identification of 558 million PSMs. Specifically, FragPipe yielded 445 million target PSMs under a 1% false discovery rate (FDR) threshold and 460 million decoy PSMs without FDR filtering. In comparison, Sage, which allows each MS/MS scan to be matched with up to 10 potential decoy sequences, identified 363 million target PSMs (FDR *<* 1%) and ~ 2.9 billion decoy PSMs (unfiltered).

**Figure 1.**
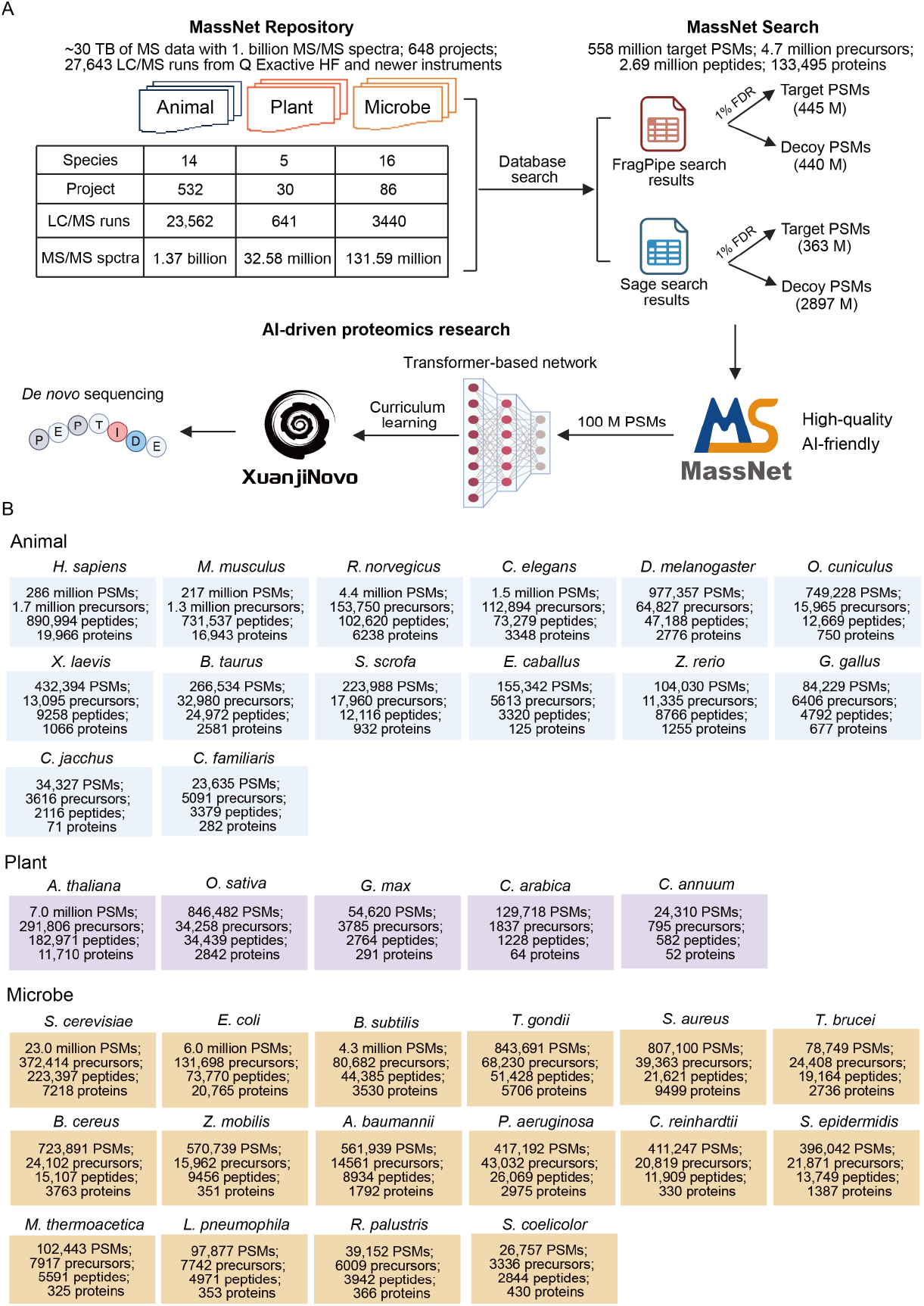
Overview of the MassNet. (A) Schematic representation of the MassNet repository architecture and data processing workflow. MassNet comprises 27,643 DDA-MS data files spanning three major biological domains: animals, plants, and microorganisms. The dataset contains ~ 30 TB raw MS data, totaling ~ 1.5 billion MS/MS spectra across 648 individual projects. Database searching was performed using FragPipe and Sage, with identifications filtered at 1% FDR. All DDA files generated from in-house experiments were considered part of a single project. The lower panel illustrates the application of 100 million PSMs in AI-driven proteomics research, including the *de novo* sequencing model xuanjiNovo, which was developed under a curriculum learning framework. (B) Overview and taxonomic composition of the MassNet dataset: covering representative species across multiple biological domains and presenting identification results at different levels (including PSMs, precursors, peptides, and proteins) for each species-specific subset.

MassNet was developed from high-quality proteomics data spanning 35 species, including 14 animals, 5 plants, and 16 microorganisms, providing a robust and comprehensive foundation for cross-species comparative analysis and AI-driven proteomics research. The dataset includes over 5.58 million PSMs, 4.7 million peptide precursors, 2.69 million unique peptides, and 133,495 proteins (**Figure 1B**). The human subset comprises 15,657 DDA-MS files, generating more than 286 million PSMs, and covering over 1.7 million peptide precursors, ~ 891,000 unique peptide sequences, and 19,966 proteins. Notably, protein identification achieves 98% coverage of the reviewed human proteins annotated in the UniProt database. The mouse dataset contains ~ 217 million PSMs, mapping to ~ 1.3 million precursors, 731,537 unique peptides, and over 16,900 proteins. Other representative model organisms, such as *R. norvegicus, C. elegans*, and *D. melanogaster*, contributed 4.4 million, 1.5 million, and ~ 978,000 PSMs, respectively, with identified protein annotation coverage exceeding 74%. MassNet also provides high-quality proteomics resources for plants. For example, the *A. thaliana* dataset contains over 7 million high-confidence PSMs, corresponding to ~ 292,000 peptide precursors, 182,971 unique peptides, and 11,710 proteins. In addition, proteomic data from *O. sativa* (rice), *G. max* (soybean), *C. arabica* (coffee), and *C. annuum* (pepper) are included, contributing over 1 million PSMs and 3177 proteins in total, significantly expanding the phylogenetic coverage of plant species. In the microbial domain, the *S. cerevisiae* subset contains more than 23 million PSMs, spanning ~ 370,000 peptide precursors, 223,397 unique peptides, and 7218 proteins, providing valuable data for studying key biological processes in eukaryotic microbes, such as metabolic pathways, signal transduction, and protein–protein interactions. In addition, the subsets for *E. coli* and *B. subtilis* each contain more than 4 million PSMs. the dataset also includes spectral data from multiple archaea, actinomycetes, and fungi, offering a comprehensive resource for microbial proteomics research. Detailed information on the number of PSMs, precursors, and identified proteins for each species are provided in **Supplementary Table 1**.

### 2.2 MSDT format facilitates scalable and efficient data processing

Spectral data were first integrated with database search results, then transformed into a structured tensor representation, and stored in columnar format using Apache Parquet. This unified, AI-optimized data format is referred to as the Mass Spectrometry Data Tensor (MSDT) (**Figure 2A**). Each mzML file was transformed into two corresponding MSDT files, resulting in a total of 55,286 MSDT files with a combined volume of ~ 3.2 TB. Compared to the raw MS data, the MSDT format achieved an average file size reduction of ~ 90%. To assess the applicability of the MSDT format across different MS platforms, we benchmarked its performance against other common storage formats using representative datasets acquired from Orbitrap, timsTOF, and TripleTOF platfoms, with a focus on two key performance metrics: data compression ratio and data loading speed. To ensure the representativeness of the results, at least six DDA-MS files were selected from each platform. The results indicated that, on the Orbitrap platform, the csv gz format achieved the highest average compression ratio (~ 3.9 *×*), followed by parquet and feather, while the tsv format exhibited the lowest compression efficiency (**Figure 2B**). For datasets acquired using Bruker or TripleTOF instruments, both parquet and feather exhibited consistent and effective compression performance across platforms. In terms of read speed, for Orbitrap data (**Figure 2C**, left panel), feather and ipc formats showed the fastest read times (*<* 1 s), followed closely by parquet (~ 1.4 s), while csv gz showed the slowest performance (~ 12.9 s). Similar trends were observed for Bruker (**Figure 2C**, middle panel) and TripleTOF (**Figure 2C**, right panel), where ipc, feather, and parquet consistently achieved low read latencies.

**Figure 2.**
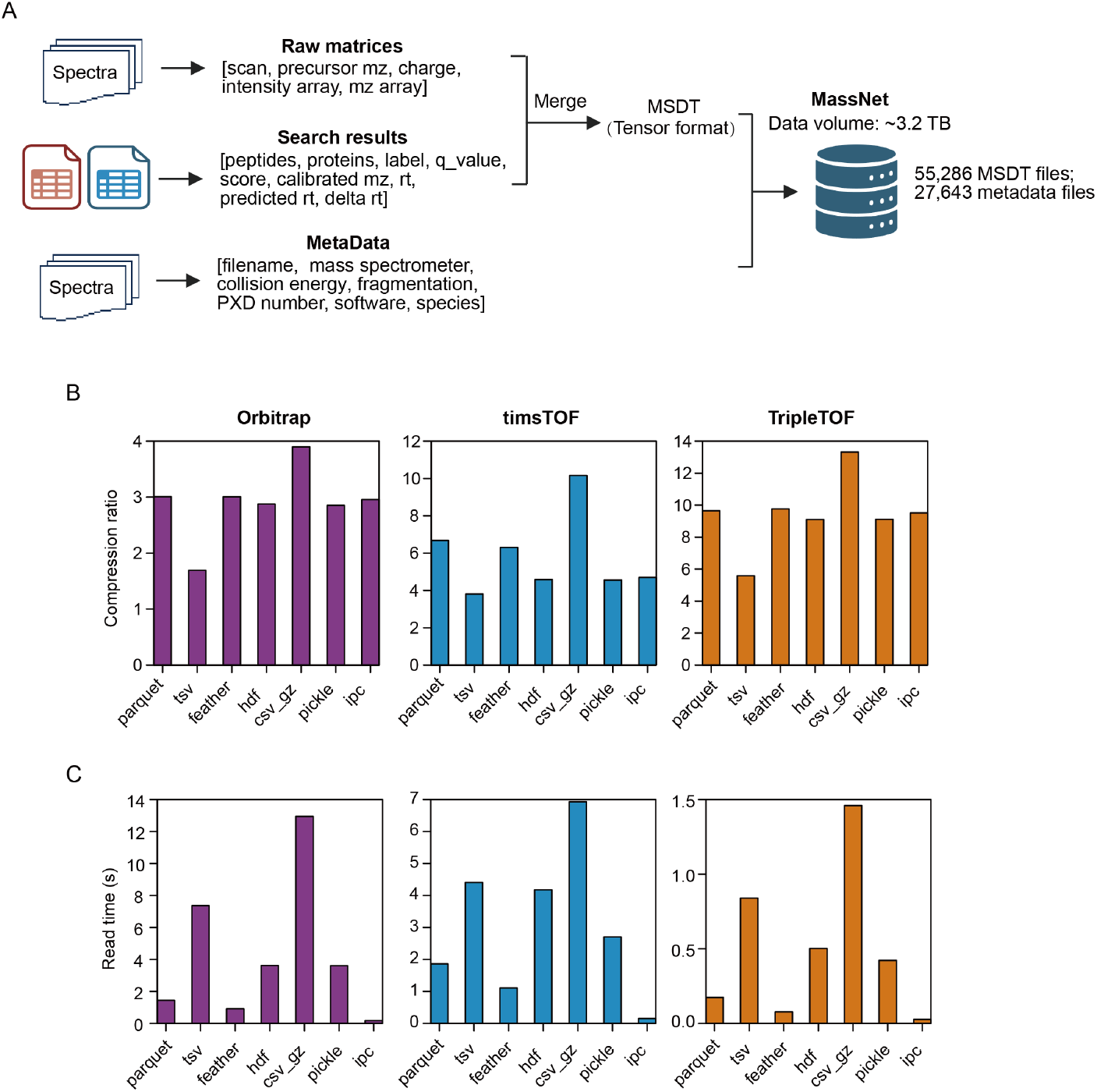
Programmatic structuring of the MassNet database and performance evaluation of the MSDT format. (A) Overview of the programmatic construction of the MassNet dataset. Raw spectral data and identification results are programmatically converted into a standardized tensor format and stored as Parquet files. MSDT files, together with the corresponding metadata, constitute the MassNet dataset, comprising 55,286 MSDT files and 27,643 metadata files, with a total size of approximately 3.2 TB. (B) Compression ratios of different serialization formats across three instrument types (Orbitrap, timsTOF, and TripleTOF). The formats tested include parquet, tsv, feather, hdf5 (hdf), csv gz (compressed), pickle, and ipc. The compression ratio is calculated by summing the original file size (.d file size for tims data, mzML file size for all other data) and the size of the search result file, then dividing by the size of the search result in each target format. (C) Read time performance (in seconds) for each format across the same instrument platforms. For each type of mass spectrometer, at least six raw files were randomly selected, and the average read time for each format was calculated.

In addition, we evaluated the generation time of various data serialization formats across the three MS platforms to assess their suitability for high-throughput data preprocessing (**Supplementary Figure 1**). The results, presented on a logarithmic scale, show that across all platforms, the ipc format consistently achieved the fastest generation time, averaging less than one second. Feather and Parquet also demonstrated strong performance. In contrast, tsv and csv gz formats required significantly longer generation times, indicating lower efficiency. The results indicate that although tsv and csv gz formats offer higher compression ratios, their relatively long generation times limit their practicality in real-time or large-scale workflows where high processing throughput is essential. In contrast, feather and parquet formats not only provide faster generation times but also maintain stable read performance, making them more suitable for high-throughput proteomics pipelines. Overall, both feather and parquet demonstrated strong performance in compression efficiency, read speed, and generation time. Given parquet’s broader ecosystem compatibility, superior support for complex data types, and widespread adoption in distributed processing frameworks such as Spark and Hadoop, it was ultimately selected as the standard storage format for MassNet to better support high-throughput and scalable proteomics data analysis.

### 2.3 Curriculum learning and non-autoregressive design enhance stability and accuracy in XuanjiNovo

To address the long-standing limitations of *de novo* peptide sequencing in terms of optimization stability and predictive accuracy, we developed XuanjiNovo, a deep learning model that integrates a non-autoregressive Transformer architecture with a progressive curriculum learning framework (**Figure 3**). In contrast to traditional autoregressive approaches that decode amino acids sequentially, XuanjiNovo generates full peptide sequences in parallel, substantially improving inference efficiency. At the heart of the model is a Transformer-based spectrum encoder that captures the fine-grained patterns of tandem mass spectrometry (MS/MS) data. Training incorporates a novel connectionist temporal classification (CTC) path sampling strategy alongside curriculum learning (**Figure 3A**), which jointly enhance optimization stability and generalization. During inference, the model first generates coarse peptide predictions that are iteratively refined, yielding substantial gains in final prediction accuracy (**Figure 3B**).

**Figure 3.**
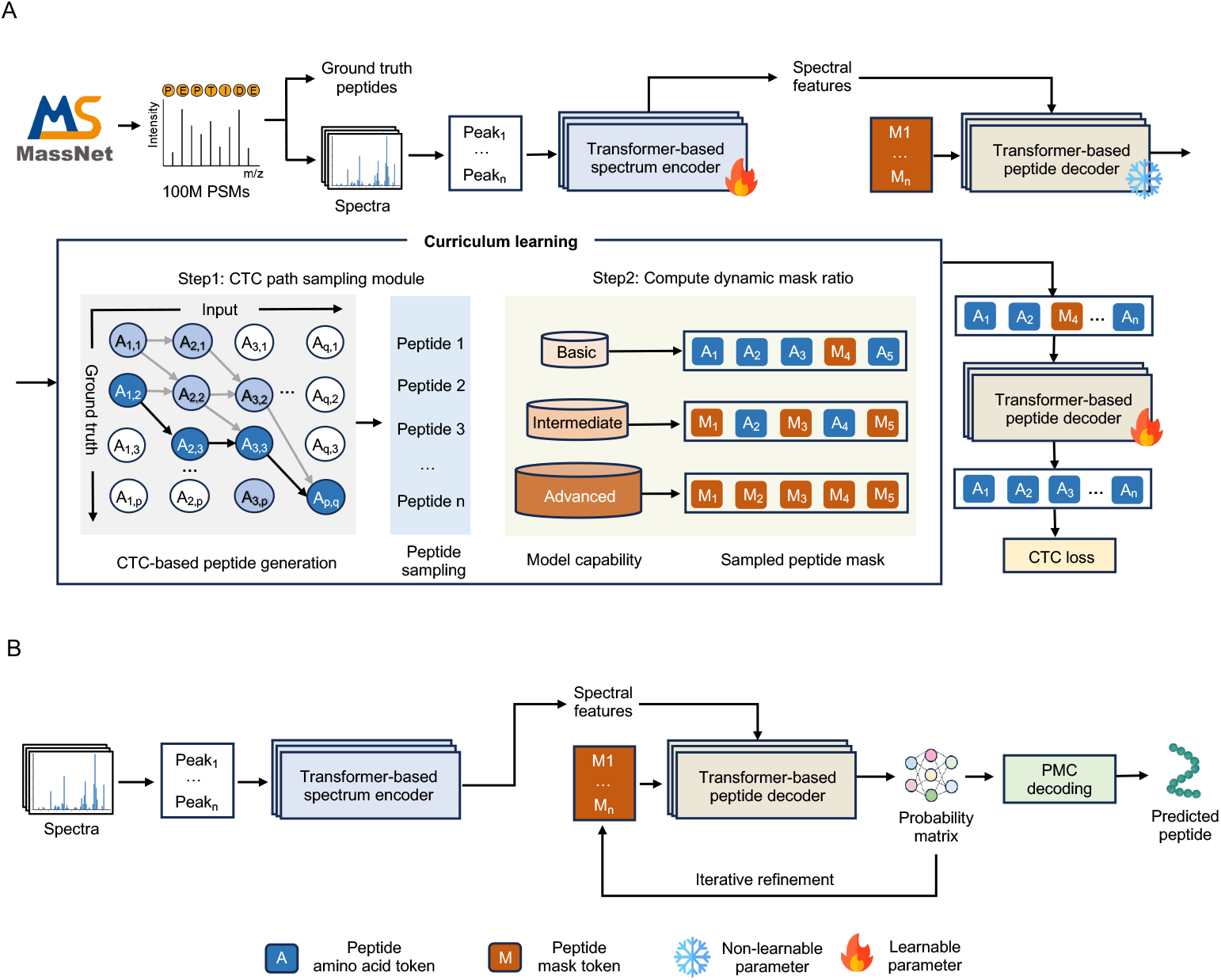
Architecture of XuanjiNovo for *de novo* peptide sequencing. (A) Training Phase: MS/MS spectra (containing peak tokens *P*_*i*_) are fed into a Transformer-based spectrum encoder. The resulting features (including blank tokens *B*_*i*_) are processed by a non-autoregressive peptide decoder. Our novel CTC path sampling module generates diverse peptide candidates (*A*_*i,j*_ representing amino acids) from ground truth sequences to improve learning robustness. Simultaneously, curriculum learning progressively enhances model capability by adjusting input mask tokens (*M*_*i*_) across basic to advanced training stages. (B) Inference Phase: For a given spectrum, features are extracted and decoded. An initial peptide sequence (*A*_1_, …, *A*_*n*_) is generated from the decoder’s probability matrix via an argmax operation. This sequence subsequently undergoes our novel iterative refinement step to boost final prediction accuracy. CTC: Connectionist Temporal Classification; PMC: Precise Mass Control.

To systematically evaluate the performance of xuanjiNovo, we constructed a 15-species benchmark dataset. Compared to the widely used 9-species benchmark dataset [3], which includes 175 files, 6.63 million MS/MS spectra, and 1.53 million PSMs, the 15-species dataset represents a substantial expansion in both scale and depth. It comprises 325 files, with the number of MS/MS spectra and PSMs increased by approximately 5-fold and 6-fold, respectively (**Supplementary Table 2**). Even when compared to another version of the 9-species dataset [36], the 15-species dataset offers 2.2 times more spectra and 3.4 times more PSMs, providing a significantly more challenging and representative benchmark for assessing model generalization, robustness, and adaptability across complex biological samples.

XuanjiNovo consistently outperformed mainstream *de novo* sequencing algorithms, including *π*-HelixNovo, CasaNovo V2, InstaNovo, ContraNovo, and PrimeNovo, across all species and at varying spectrum coverage depths (**Figure 4A**). The average peptide recall per species, as summarized in **Figure 4B**, shows that XuanjiNovo achieved peptide recall values of 0.45 on the *H. sapiens* dataset and 0.68 on the *M. musculus* dataset. These results are significantly higher than those of the second-ranked PrimeNovo (0.32 and 0.60) and the third-ranked InstaNovo (0.26 and 0.52), respectively. Notably, the *H. sapiens* dataset was acquired using the latest Orbitrap Astral MS platform, indicating XuanjiNovo’s strong adaptability to high-sensitivity, high-resolution instrumentation. On the particularly challenging *S. lycopersicum* dataset, XuanjiNovo still achieved a recall of 0.25, significantly outperforming all other models, whose performance ranged from 0.001 (HelixNovo) to 0.06 (ContraNovo). These results highlight XuanjiNovo’s remarkable robustness under low-coverage and noisy conditions. For *S. cerevisiae*, XuanjiNovo achieved a peptide recall of 0.69, comparable to PrimeNovo and substantially higher than other methods.

**Figure 4.**
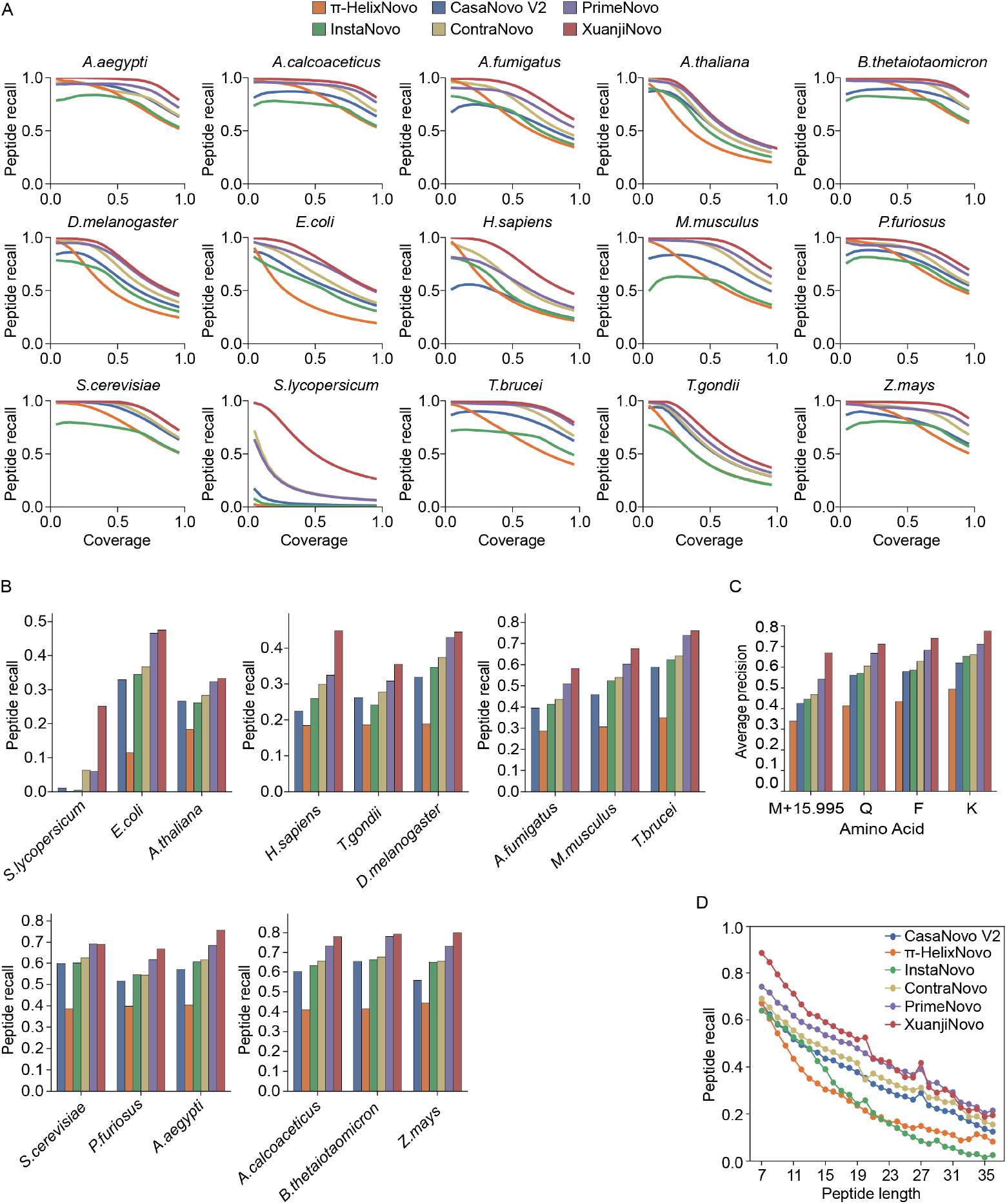
Comparison of XuanjiNovo and existing *de novo* peptide sequencing models on the 15-species benchmark dataset. (A) Comparison of peptide recall versus coverage across 15 diverse species for six models: *π*-HelixNovo, InstaNovo, CasaNovo V2, ContraNovo, PrimeNovo, and XuanjiNovo. (B) Cross-species peptide recall aggregated by species, illustrating overall performance differences across taxa. (C) Average identification accuracy of near-isobaric amino acid residues across different models, including lysine (K, 128.094963 Da) vs. glutamine (Q, 128.058578 Da) and phenylalanine (F, 147.068414 Da) vs. oxidized methionine (M+15.995, 147.035400 Da). (D) Peptide recall across different peptide lengths is compared among six models: HelixNovo, CasaNovo V2, InstaNovo, ContraNovo, PrimeNovo, and XuanjiNovo. Unless otherwise stated, xuanjiNovo in this study refers to xuanjiNovo-100M. The *π*-HelixNovo model is publicly available and was trained on datasets derived from eight species, excluding human.

One of the key challenges in *de novo* sequencing is accurately distinguishing amino acids with near-isotopic masses. Compared to models such as PrimeNovo, XuanjiNovo exhibited superior discrimination in identifying these closely related amino acid pairs (**Figure 4C**). For example, in differentiating lysine (K, 128.094963 Da) from glutamine (Q, 128.058578 Da), XuanjiNovo achieved recognition accuracies of 0.78 and 0.71, respectively. Similarly, for phenylalanine (F, 147.068414 Da) and oxidized methionine (M, 147.035400 Da), the model also achieved significantly higher precision than other methods.

In addition, we evaluated the predictive performance of XuanjiNovo across different peptide lengths (**Figure 4D**). The results showed that XuanjiNovo consistently outperformed all other models, with particularly strong performance on peptides shorter than 24 amino acids. These findings further highlight XuanjiNovo’s strong generalization capability and broad applicability. The model delivers stable, high-quality predictions across diverse biological contexts, making it a powerful tool for proteomic analysis in complex and heterogeneous MS datasets.

### 2.4 Scaling training data unlocks superior model convergence, predictive accuracy, and crossspecies generalization

To assess the impact of training data size on the performance of XuanjiNovo, we trained the model using three subsets of the MassNet dataset, containing 30 million (XuanjiNovo-30M), 65 million (XuanjiNovo-65M), and 100 million PSMs (XuanjiNovo-100M), respectively. As shown in **Figure 5**, we compared the performance of XuanjiNovo and PrimeNovo on a validation set derived from MassNet. This validation set was carefully constructed to ensure that its peptide sequences were entirely independent of those in the training set, providing a rigorous and objective assessment of model performance. Compared with XuanjiNovo-30M and XuanjiNovo-65M, XuanjiNovo-100M exhibited faster convergence and greater training stability, maintaining the lowest validation loss throughout the training process (**Figure 5A**). This superior learning process ultimately results in improved predictive performance. In terms of amino acid-level accuracy (**Figure 5B**), XuanjiNovo-100M achieved an accuracy of approximately 0.82, outperforming both XuanjiNovo-65M (0.78) and XuanjiNovo-30M (0.75). Notably, this represents a *>* 20% improvement over the established PrimeNovo model, which achieves around 0.60. A consistent performance gain was also observed in peptide-level recall (**Figure 5C**), where XuanjiNovo-100M reached ~ 0.68, an increase of ~ 14% compared to the 30M model (~ 0.54), and substantially higher than PrimeNovo’s recall of ~ 0.30, highlighting a significant enhancement in peptide identification sensitivity. Next, we evaluated the effect of training set size on XuanjiNovo’s cross-species generalization using a challenging benchmark dataset comprising 15 phylogenetically diverse species. As shown in **Figure 5D**, XuanjiNovo-100M consistently achieved the highest peptide recall rates across all evaluated species. For example, in *E. coli*, recall improved from 0.35 (XuanjiNovo-30M) to 0.48 (XuanjiNovo-100M); in *A. thaliana*, from 0.29 to 0.33 (a 4% increase); and in *D. melanogaster*, from 0.37 to 0.44 (a 7% increase). Notably, performance on the human dataset exhibited the greatest sensitivity to training set size, with peptide recall increasing markedly from 0.36 to 0.45, a nearly 10% improvement. Even in species where strong performance was already achieved with the 30M dataset, such as *B. thetaiotaomicron* and *Z. mays*, both exceeding a peptide recall of 0.72, increasing the training set to 100M further improved recall to 0.79 and 0.80, respectively, highlighting that large-scale training remains beneficial even under high baseline performance conditions. These results highlight the critical role of large-scale, high-quality datasets in enhancing the accuracy, robustness, and cross-species generalizability of *de novo* peptide sequencing models, establishing a key strategy for advancing their application in broader biological research.

**Figure 5.**
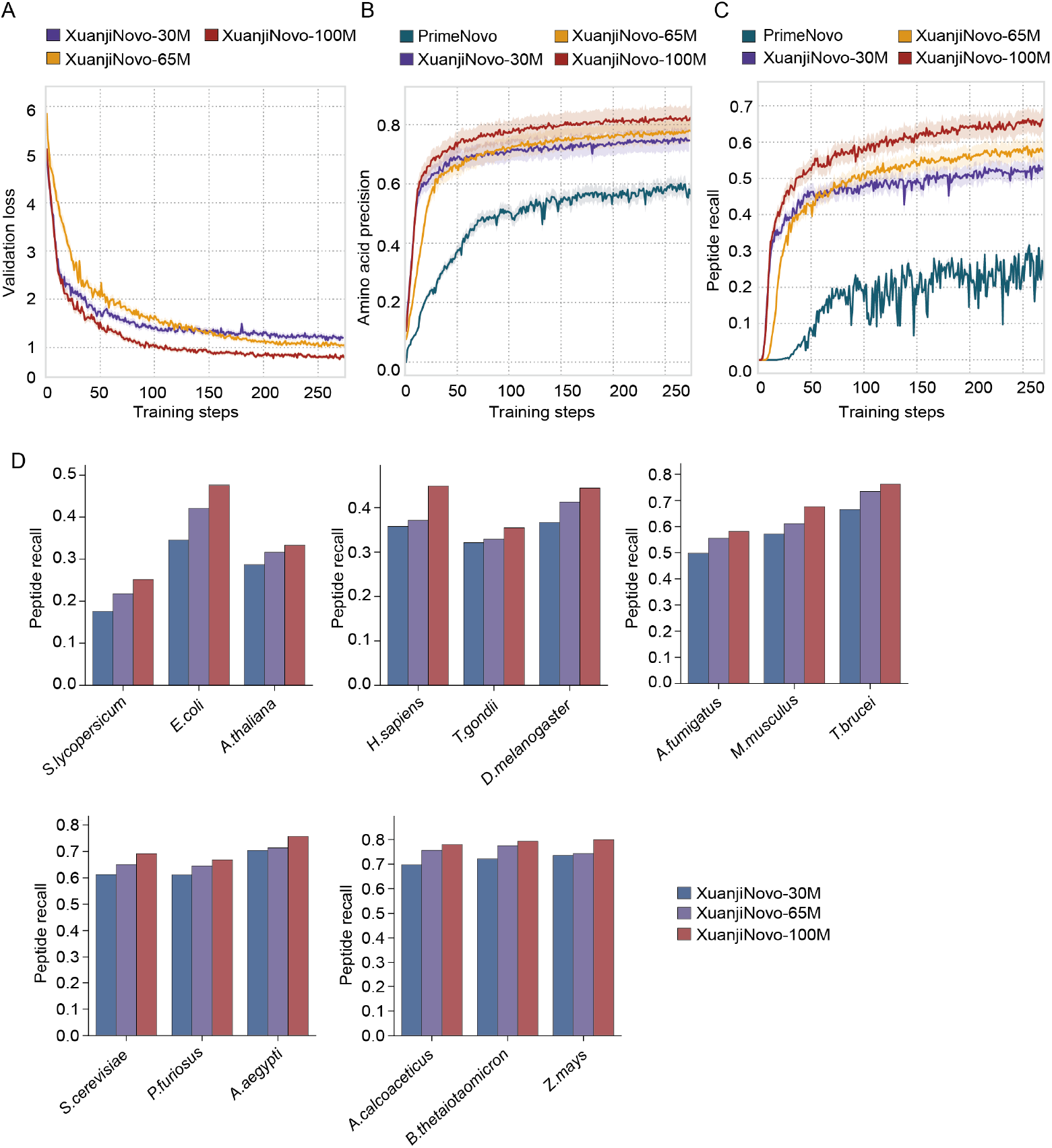
Comparison of model performance with training sets of 30 million (XuanjiNovo-30M), 65 million (XuanjiNovo-65M), and 100 million PSMs (XuanjiNovo-100M). (A–C) Training curves of XuanjiNovo models with different dataset sizes (30M, 65M, 100M) compared to PrimeNovo: (A) validation loss, (B) amino acid precision, and (C) peptide recall. Direct comparison of loss values with PrimeNovo is not applicable due to differing loss functions. (D) Peptide recall of XuanjiNovo models evaluated on the 15-species benchmark at all three dataset scales (30M, 65M, 100M).

### 2.5 XuanjiNovo achieves robust generalization across diverse datasets and MS platforms through fine-tuning

Technical heterogeneity across MS platforms remains a major challenge to achieving robust model generalization. To systematically evaluate the generalization performance of XuanjiNovo under cross-dataset and cross-platform scenarios, we compared its performance against five state-of-the-art *de novo* peptide sequencing models, including *π*-HelixNovo, InstaNovo, ContraNovo, Casanovo V2, and PrimeNovo, on three representative benchmark datasets. The results showed that on the nine-species benchmark dataset, XuanjiNovo-100M exhibited lower peptide recall rates than existing models across most species, with only marginal advantages observed in the human and mouse datasets (**Supplementary Figure 2A**), indicating that cross-platform robustness remains a significant challenge. To address the challenges of cross-platform variability and enhance overall model robustness, we implemented a targeted fine-tuning strategy. Specifically, we further trained the XuanjiNovo-130M model using 30 million PSMs from the MassIVE-KB dataset to improve its adaptability across heterogeneous data sources. As expected, the XuanjiNovo-130M model demonstrated substantial performance improvements across all evaluated species. The most significant gains were observed in *S. cerevisiae* and *B*.*subtilis*, where peptide recall increased by 64.6% and 60.2%, respectively. Even in the human dataset, which showed the smallest improvement, XuanjiNovo-130M achieved a 1.9% performance gain, reaching a peptide recall of 0.60. Although slightly lower than ContraNovo (0.62), it still outperformed all other existing models. Notably, in the *M. musculus* dataset, XuanjiNovo-130M achieved the highest peptide recall, reaching approximately 0.67. In other representative species such as *C. Cloacimonas, M. mazei*, and *S. cerevisiae*, the model maintained strong competitiveness, ranking second in peptide recall. These results indicate that the fine-tuned XuanjiNovo has substantially improved its cross-platform adaptability, effectively overcoming the generalization challenges observed in earlier versions and demonstrating enhanced robustness and broader applicability.

In addition to the 9-species benchmark dataset, we further assessed the generalization capability of XuanjiNovo using two independent datasets acquired on distinct MS platforms. For the HCC dataset [37], which consists of human hepatocellular carcinoma tissue samples acquired using the Orbitrap Fusion platform, XuanjiNovo-100M achieved a peptide recall rate of 0.68 without any fine-tuning, substantially outperforming the next best model, PrimeNovo (0.38), with an improvement of ~ 77% (**Supplementary Figure 2B**). Similarly, on the PT dataset [38], which includes synthetically generated peptides acquired using the Orbitrap Fusion Lumos platform, XuanjiNovo-100M achieved a recall of 0.62 without fine-tuning, outperforming existing models by margins ranging from 3.2% to 31.9 (**Supplementary Figure 2C**). These findings highlight XuanjiNovo’s strong generalization across diverse biological sample types and experimental conditions, with particularly notable performance on high-resolution MS platforms. They also emphasize the importance of continuous model refinement to effectively accommodate heterogeneous and multi-source data environments.

## 3 Discussion

MassNet represents a critical milestone in the integration of AI and proteomics, comparable in significance to WordNet and OpenWebText in natural language processing and ImageNet in computer vision. It is the first AI-friendly MS dataset that combines both scale and standardized structure, specifically designed to support deep learning-based proteomic analysis. MassNet comprises approximately 28,000 DDA files and nearly 30 TB of raw MS data, it contains over 1.5 billion MS/MS spectra, spanning 35 biological species, including animals, plants, and microorganisms, providing a comprehensive spectral representation across diverse biological taxa. Notably, the human subset alone contains 286 million PSMs and 1.7 million precursors, corresponding to 19,966 proteins, which accounts for 97.7% of all reviewed human proteins annotated in UniProt. This dataset not only establishes a solid foundation for large-scale training of deep learning models in proteomics but also provides a unified and generalizable data framework suitable for multi-species and cross-platform proteomic research.

A key innovation of MassNet lies in its AI-oriented design, which offers scalable, structured, and highthroughput access to MS data, facilitating advanced machine learning applications. Although public repositories have accumulated vast amounts of MS data, much of it remains underutilized due to inconsistent data formats and incomplete metadata [21]. For example, MassIVE-KB [27], a subproject of MassIVE, integrates 27,404 DDA files to construct a high-confidence, peptide-centric human proteome knowledge base. While it plays a vital role in improving proteome coverage and reidentification performance, its primary focus remains on optimizing protein identification within traditional proteomics workflows, rather than on developing data structures specifically tailored for AI applications. In contrast, MassNet is built from the ground up with AI compatibility as a foundational principle, aiming to bridge the gap between raw MS data and deep learning applications. Commonly used MS data formats such as .mzML and .mgf require disk access prior to parsing, leading to frequent and costly I/O operations. These files are typically large, ranging from hundreds of megabytes to several gigabytes, which further increases storage requirements and access overhead. Such limitations pose significant barriers to scalable data processing and markedly compromise the overall efficiency of analysis [39]. MassNet adopts a unified data representation strategy by structuring raw spectra into two-dimensional tensors that preserve key features such as mass-to-charge ratio (m/z) and intensity. All data are stored in Apache Parquet format, which offers advantages such as columnar storage, compression, and schema definition. This format enables efficient batch reading and integrates seamlessly with the DataLoader modules of mainstream deep learning frameworks, including PyTorch [40] and TensorFlow [41], thereby streamlining data preprocessing. The ready-to-use format not only significantly accelerates model training on hardware accelerators such as GPUs and TPUs but also improves training reproducibility. By lowering the technical barrier for AI researchers and developers working with MS data, it substantially enhances the usability and scalability of deep learning methods in proteomics analysis. In addition, each spectrum is paired with comprehensive metadata, including instrument type, fragmentation method, sample origin, and corresponding PSM information, facilitating multi-dimensional and multi-scale modeling and analysis. MassNet also emphasizes broad coverage and diversity to ensure robustness and generalizability across different experimental conditions, tissue types, and species.

*De novo* peptide sequencing enables the direct inference of peptide sequences from MS/MS spectra without relying on reference databases, making it essential for the analysis of novel proteins, non-model organisms, and unannotated regions of the proteome. While recent deep learning–based approaches, such as DeepNovo [3], SM-SNet [42], pNovo 3 [43], RANovo [44], PointNovo [45], DEPS [46], DPST [47], Casanovo [48], PepNet [49], SeqNovo [50], Denovo-GCN [51], BiATNovo [52], GraphNovo [53], PGPointNovo [54], *π*-HelixNovo [5], NovoB [55], Casanovo V2 [4], AdaNovo [56], *π*-xNovo[57], ContraNovo [58], Spectralis [59], RankNovo [12], InstaNovo [6], and PrimeNovo [7] have shown promising performance on benchmark datasets, they often face challenges when applied to large-scale, diverse datasets. Issues such as training instability, slow inference speed, and limited generalizability hinder their effectiveness and constrain their broader applicability in real-world proteomics. To address this, we propose a robust curriculum learning paradigm [31], in which the model initially predicts only partial peptide sequences, with the remaining labels provided as contextual guidance. This approach reduces early-stage learning difficulty and prevents training collapse in complex or imbalanced datasets. As training progresses, task difficulty is gradually increased based on model performance, enabling smooth and effective convergence. The model adopts a non-autoregressive Transformer architecture [32] that supports bidirectional peptide sequence modeling and significantly accelerates inference. Combined with sequence-level self-refinement and a precise mass control (PMC) module, the model is trained on 100 million spectra from MassNet and consistently outperforms existing *de novo* sequencing methods across multiple tasks. While the model’s performance on the widely used nine-species benchmark dataset appears less competitive, this is likely due to differences in instrumentation. Specifically, the benchmark data are acquired using earlier-generation Q Exactive instruments, whereas XuanjiNovo was trained on spectra generated from Q Exactive HF or newer high-resolution MS platforms. Such cross-platform differences may lead to significant variations in spectral signal distributions, increasing the uncertainty and complexity of model generalization across instrumentation types. Following a simple fine-tuning step using 30 million PSMs from the MassIVE-KB dataset, XuanjiNovo showed notable performance improvement. This enhancement further supports the observation that MassIVE-KB contains substantial data acquired from earlier instruments (e.g., Q Exactive), which likely improves the model’s adaptability to legacy platforms. Overall, the superior performance of XuanjiNovo compared to existing models is largely attributed to the unprecedented scale and cross-species diversity of the MassNet dataset. This extensive resource enables the model to effectively learn complex fragmentation patterns and contextual sequence dependencies, thereby enhancing its generalizability and real-world applicability.

MassNet sets a new benchmark for AI-optimized MS infrastructure by delivering a well-annotated, standardized, and scalable dataset. It can be potentially applied to a wide spectrum of proteomics tasks, including peptide identification, PTM prediction, *de novo* sequencing, and protein quantification. By overcoming persistent data bottlenecks in AI-driven proteomics, MassNet establishes a robust foundation for the next generation of data-centric modeling paradigms in computational biology. Through standardized data representation, democratized access, and large-scale machine learning deployment at an unprecedented scale, MassNet is positioned to accelerate both innovation and reproducibility in computational proteomics.

## 4 Materials and Methods

### 4.1 MassNet dataset

The MassNet dataset comprises ~ 30 TB of high-quality MS data, including over 1.5 billion MS/MS spectra from 648 projects and 27,643 LC/MS runs generated using Q Exactive HF and newer instruments. As shown in **Figure 1**, the dataset spans a broad range of biological domains, covering 14 animal species (532 projects, 23,562 runs, 1.37 billion spectra), 5 plant species (30 projects, 641 runs, 32.58 million spectra), and 16 microbial species (86 projects, 3,440 runs, 131.59 million spectra). To construct MassNet, raw MS/MS files were first converted to mzML format using msconvert GUI (ProteoWizard, v3.0.19014), except for the .d format files generated by trapped ion mobility spectrometry (TIMS), which can be used directly for searching without conversion. Database searches were then performed using Sage (v0.14.6) [35] and FragPipe (v21.1) [9, 34]. All species, except for *T. gondii* and *T. brucei*, used unreviewed protein sequences from TrENBL, while the remaining species used reviewed sequences. All FASTA files were downloaded from the UniProt database. The search parameters included: peptide length of 7-50, mass tolerance of 20 ppm, trypsin digestion (up to two missed cleavages), variable modifications (methionine oxidation and N-terminal acetylation), and fixed carbamidomethyl cysteine modification. FragPipe includes MSBooster, Percolator, and Philosopher for the downstream processing of MSFragger search results. For Sage, each fragment spectrum yields up to 10 candidate peptide sequences. The PSM file (“psm.tsv”), precursor ion file (“ion.tsv”), peptide file (“peptide.tsv”), and protein file (“protein.tsv”) generated by FragPipe were used for downstream analysis. The “spectrum q”, “peptide q”, and “protein q” fields in the search result file (“results.sage.tsv”) produced by the Sage represent the q values at the PSM, peptide, and protein levels, respectively, and were used as criteria for subsequent filtering. Following spectrum q-based filtering, precursor ions were deduplicated to generate the final precursor ion set. All results were filtered using a q value threshold of 0.01 to ensure high-confidence identifications.

### 4.2 Mass spectrometry DDA tensor (MSDT)

For each file, spectral data were extracted from the corresponding mzML files, capturing key raw matrix components such as scan number, precursor m/z, charge state, intensity array, and m/z array. These raw spectral matrices were then integrated with search engine outputs from FragPipe or Sage, which provided peptide and protein identifications along with additional attributes of candidate peptides, including peptide sequences, protein accessions, identification scores, q-values at various levels (PSM, peptide, and protein), calibrated m/z values, retention time (RT), predicted RT, and delta RT. The raw spectral matrices and identification results are programmatically integrated into a standardized tensor format, stored efficiently in Parquet files and referred to as MSDT. Each MSDT file encapsulates both spectral feature vectors and identification annotations in an AI-friendly structure, facilitating direct access for model training and inference. In addition, structured metadata files were compiled to comprehensively document instrument configurations, acquisition parameters, and analysis workflows. These include details such as the mass spectrometer model, fragmentation method, collision energy, PXD identifier, software name and version, and species information. This rich metadata forms a complete and traceable framework from data acquisition to interpretation, ensuring both dataset traceability and reproducibility.

To evaluate the performance of the MSDT format, MS data acquired from three types of mass spectrometers, including Orbitrap, timsTOF, and TripleTOF, were used. The Orbitrap data were obtained using the Orbitrap Astral instrument and are available from the PRIDE repository under the identifier PXD046453 [60]. The Bruker dataset consisted of eight samples used for spectral library construction, all generated in-house with the timsTOF Pro mass spectrometer. The TripleTOF dataset, also produced in-house, included ten DDA files acquired using the SCIEX TripleTOF 6600 instrument in IDA mode. Except for the Bruker .tims raw files, all datasets were converted to the mzML format using a unified processing pipeline. All raw data conversions and preprocessing steps followed standardized protocols to ensure consistency and fairness.

### 4.3 15-species benchmark dataset

To construct a representative and high-quality benchmark dataset for *de novo* peptide sequencing, we curated and processed a total of 325 high-quality DDA files derived from 15 representative species across multiple biological kingdoms. The MS data were acquired using three high-resolution MS platforms: Q Exactive HF, Orbitrap Exploris 480, and Orbitrap Astral, ensuring both spectral diversity and quality across instrument types. The benchmark dataset spans a taxonomically diverse selection of species, organized as follows: Animals: *Homo sapiens, Mus musculus, Aedes aegypti*, and *Drosophila melanogaster*; Plants: *Arabidopsis thaliana, Zea mays*, and *Solanum lycopersicum*; Microbe: *Escherichia coli, Saccharomyces cerevisiae, Pyrococcus furiosus, Aspergillus fumigatus, Toxoplasma gondii, Trypanosoma brucei, Bacteroides thetaiotaomicron*, and *Acinetobacter calcoaceticus*. All raw MS data were processed using a standardized preprocessing pipeline, consistent with that applied in the construction of MassNet. Database searches were conducted on these benchmark datasets using FragPipe (v21.1) [9, 34], and identification results were filtered at a 1% FDR threshold. Detailed information on species distribution and project accession numbers is provided in **Supplementary table 2**.

### 4.4 Training datasets

We developed XuanjiNovo, a *de novo* peptide sequencing model designed for large-scale, heterogeneous DDA-MS data. For model training, 100 million high-confidence PSMs were randomly sampled from the MassNet dataset. These PSMs span multiple species, MS platforms, fragmentation methods (e.g., HCD, CID), and experimental conditions, ensuring dataset diversity and enhancing the model’s generalization capability. During training, XuanjiNovo receives input in the form of preprocessed feature tensors formatted in the standardized MSDT structure, which encapsulates both spectral features and their corresponding peptide identifications. To effectively manage the scale and complexity of this dataset, XuanjiNovo adopts a non-autoregressive Transformer backbone, inspired by the success of earlier models such as PrimeNovo [7]. A detailed description of the model architecture is provided in the following section.

### 4.5 Formulation

The input to our model is a set of peaks extracted from a tandem mass spectrum, formally represented as:

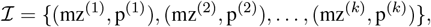

where each pair consists of a mass-to-charge ratio value mz^(*i*)^ and its corresponding intensity p^(*i*)^. Along with the peak set, the model also receives two scalar values: the precursor mass *m ∈* ℝ, representing the total mass of the peptide ion prior to fragmentation, and the charge state *z ∈* ℕ of the ion.

The output is a sequence of amino acids 𝒜 = (*a*_1_, *a*_2_, …, *a*_*n*_) that best explains the input spectral data.

The prediction task is framed as a sequence generation problem conditioned on the input spectrum, such that the generated peptide aligns with the fragmentation captured by *ℐ* and satisfies the global constraint imposed by *m*.

### 4.6 Model Architecture

Our model leverages a non-autoregressive Transformer (NAT) architecture [32], which has been shown highly effective for biological sequence modeling [7], consisting of a spectrum encoder, a NAT peptide decoder, and a final decoding module that enforces precise mass constrain (PMC). The overall design supports curriculum learning and enables iterative refinement during inference.

#### Spectrum Encoder

The encoder transforms the spectrum ℐ into a dense, informative latent representation suitable for peptide sequence generation. Each mz^(*i*)^ is embedded into a *d*-dimensional vector using sinusoidal positional encoding, which ensures that nearby masses have similar representations and allows the model to capture mass-scale relationships. The sinusoidal encoding is computed as follows:

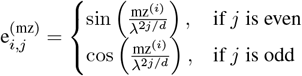

for dimensions *j* = 1, …, *d*, where *λ* is a scaling constant. The intensity p^(*i*)^ is similarly encoded into *d* dimensions and added to the corresponding mz embedding, yielding the combined input vector 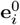 for the *i*-th peak.

The resulting sequence 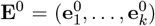 is passed through *m* Transformer encoder layers, each applying multi-head self-attention and feedforward transformations. This produces a final context-aware spectrum representation **E**^(*m*)^ *∈* ℝ^*k×d*^, which serves as input for the decoder.

#### Peptide Decoder and CTC Loss

In contrast to autoregressive models, which generate one token at a time in a left-to-right fashion, our NAT decoder predicts token probabilities at all positions simultaneously with bidirectional information to its surroundings through self-attention modules. Each position *t* in the output sequence receives an input embedding 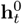 that may be initialized with either a learned start-of-sequence token, a mask token, or a partially revealed ground-truth token (detailed in curriculum learning section).

These embeddings are updated through *L* decoder layers, each performing both bi-directional self-attention among all decoding positions to access its surrounding generational information, and cross-attention with the spectrum representation **E**^(*m*)^ to match with input signals. The final hidden state 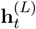 at each position is passed through a linear transformation followed by a softmax function to produce a probability distribution over the amino acid vocabulary:

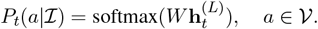

To train the model, we use the *Connectionist Temporal Classification* (CTC) loss, following its successful application in PrimeNovo. CTC is especially well-suited for tasks where the input and output sequences are unaligned—such as *de novo* peptide sequencing—because it allows the model to predict sequences without requiring exact position-wise alignment between input and target.

At the heart of CTC is the idea of alignment paths. A path **y** = (*y*_1_, *y*_2_, …, *y*_*T*_) is a length-*T* sequence of token predictions, where each *y*_*t*_ is chosen from the extended vocabulary that includes a special blank token *ϵ*. Many different paths can correspond to the same final label sequence 𝒜after being collapsed by a function Γ(·), which removes consecutive duplicates and blanks. For example,

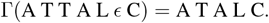

This means that the label sequence A T A L C can arise from multiple distinct prediction paths.

The CTC loss is defined to encourage the model to assign high total probability to all paths that collapse to the correct label sequence:

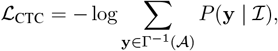

where 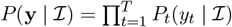 is the probability of the path **y** under the model’s output distribution.

This formulation provides several advantages: rather than requiring the model to produce one specific alignment, CTC allows it to explore multiple valid prediction paths that lead to the correct output. This flexibility enables the model to learn robustly in the presence of uncertainty or noise in token positions. In the context of peptide sequencing, where fragment peaks may be missing or imprecise, CTC allows the model to focus on producing the correct amino acid sequence regardless of exact temporal alignment, improving both accuracy and convergence.

#### Mass Constrained Decoding

To ensure preciseness of the decoding, the predicted peptide must satisfy the precursor mass constraint given by precursor mass *m*. After obtaining the position-wise probabilities *P*_*t*_(*a*|*ℐ*) for all amino acids *a*, we apply a post-processing step that selects the most probable sequence under a mass budget.

This is modeled as a variant of the 0-1 knapsack problem. Each candidate token *a* has an associated mass *M* (*a*) and a log-probability score log *P*_*t*_(*a*). The decoding process searches for a sequence (*a*_1_, …, *a*_*n*_) such that:

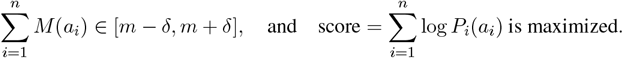

Where *δ* is the error threshold. This constrained optimization is efficiently solved via dynamic programming.

### 4.7 CTC-based Curriculum Learning for Ultra-Robust Peptide Learning

Deep learning models often struggle to adapt to changes in data distribution [61], particularly when the new distribution introduces rare or complex patterns that are hard to capture. These challenges frequently lead to optimization failures, especially in early training stages [62]. Non-autoregressive (NAT) models are particularly vulnerable to this issue: in our experiments, up to 80% of NAT model training runs failed to converge when applied to large-scale, heterogeneous proteomics datasets. This high failure rate stems from the fundamental design of NAT models [63]. Unlike autoregressive models—which predict each token *x*_*t*_ conditioned on all previous outputs *x*_1:*t−*1_, thereby narrowing the hypothesis space incrementally—NAT models attempt to predict all tokens in parallel, treating each *x*_*t*_ as independent. This independence expands the search space exponentially and severely limits the model’s capacity to capture global dependencies during training.

To alleviate these issues, we introduce a curriculum learning mechanism tailored for CTC-based NAT peptide generation. The core idea is to lower the effective difficulty of the learning task by providing partial supervision through carefully masked input sequences. Early in training, the model is guided using easy prediction targets that reveal most of the correct amino acid labels. As training progresses, the model gradually receives less supervision, learning to generalize from partial evidence.

Let **y**^*′*^ denote a selected CTC decoding path such that Γ(**y**^*′*^) = 𝒜. We define a masking operator *ρ*(𝒜, **y**^*′*^) that replaces a fraction of the tokens in **y**^*′*^ with a special [mask] token. This masked sequence is then passed as input to the decoder. The model is trained to reconstruct the full target sequence 𝒜, optimizing the following\ objective:

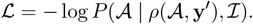

This conditioning mechanism changes the optimization from blind sequence prediction to a conditioned generation task. When *ρ*_ratio_ = 0, i.e., no masking is applied and all answer amino acids are revealed, the model sees the full ground truth and the prediction task becomes trivial. As *ρ*_ratio_ increases toward 1, the model must predict more tokens as less answer tokens are provided, thus gradually increasing difficulty. This transition can be summarized as a reduction in the effective search space:

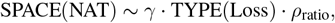

where TYPE(CTC) is considerably larger than TYPE(CrossEntropy) because each label corresponds to 𝒪 (*nT*) alignment paths rather than one. This framing provides a mathematical justification for the stability benefits observed with curriculum training.

In practice, CTC introduces a significant complication: the token positions in **y**^*′*^ do not correspond one-to-one with those in *𝒜*. This makes it impossible to simply mask position *t* in *𝒜* and expect the model to interpret it correctly. To circumvent this, we define our curriculum on **y**^*′*^ rather than *𝒜*. The detailed procedure is as follows:

**Step 1: Forward Pass to Select CTC Path**. We begin with an empty decoder input and perform a forward pass to compute *P*_*t*_(*·*|*ℐ*) at each position *t* = 1, …, *T*. From this, we enumerate valid CTC paths **y** such that Γ(**y**) = 𝒜. Each path is scored by its total probability *P* (**y** | **ℐ**) = Π_*t*_ *P*_*t*_(*y*_*t*_ |*ℐ*). We select the most probable alignment path **y**^*′*^ from this set. This process is visualized in **Figure 3A**.

**Step 2: Masking Strategy**. Given **y**^*′*^, we randomly replace a subset of its tokens with [mask]. The fraction of tokens replaced is dynamically controlled based on training progress. For example, if **y**^*′*^ = ACTTC, we may generate *ρ*(𝒜, **y**^*′*^) = A [mask] T [mask] C. This masked sequence serves as the decoder input for the current training step. This process is visualized in **Figure 3A**.

**Step 3: Training Objective**. Despite conditioning on the masked version of **y**^*′*^, the model is still trained using the original CTC objective. That is, it must maximize the total probability over *all* valid paths **y** such that Γ(**y**) = 𝒜. This encourages the model to learn the correct mapping from masked sequences to complete target alignments without collapsing to memorization.

This process enables the model to learn alignment patterns by partially observing plausible decoding paths, without ever requiring token-level supervision. It also ensures that the model generalizes over the space of all valid paths rather than overfitting to a single alignment.

**Illustrative Example**. Consider a true label 𝒜= ACTC. A few valid CTC paths might include: (1) A C T T C (2) A C T C C (3) A C *ϵ* T C. Suppose the model selects **y**^*′*^ = ACTTC. We then randomly apply masking to get *ρ*(*𝒜*, **y**^*′*^) = A [mask] T [mask] C. This input is passed through the decoder, which is trained to predict the original label *𝒜* under the CTC loss.

The partially masked path introduces two key learning signals. First, it exposes the model to likely alignment structures **y**^*′*^, which implicitly guide the attention mechanism. Second, it provides partial label information that constrains the prediction space and improves convergence. As training progresses, more tokens are masked, transitioning the model from supervised alignment recovery to full unsupervised generation.

This curriculum-guided conditioning is one of the core contributions of our model design. It transforms NAT training from a brute-force alignment discovery process into a structured, staged learning problem where difficulty increases only as the model’s competency improves. This makes our approach well-suited to diverse peptide datasets, including those with rare motifs, sparse peak coverage, or high levels of noise.

### 4.8 Inference via Iterative Curriculum Refinement

During inference, the ground-truth peptide 𝒜 is unavailable, and thus the curriculum conditioning mechanism must be adapted. To exploit the decoder’s capacity to leverage conditional inputs, we propose an iterative refinement strategy.

At the initial iteration (*i* = 0), the decoder is provided with all mask tokens. The resulting prediction **y**^(0)^ is obtained via argmax over each position’s output distribution. This predicted path is then used as conditional input for the next iteration:

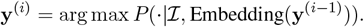

This process is repeated for *N* iterations, allowing the decoder to progressively refine its prediction. The final output **y**^(*N*)^ is then passed to the mass-constrained decoding module to ensure consistency with the precursor mass. This process is visualized in **Figure 3B**. Unlike autoregressive beam search or greedy decoding, this iterative method allows for full-sequence correction and enables the model to benefit from previously learned curriculum patterns even during testing.

## 5 Data availability

The nine-species benchmark dataset was downloaded from the Mass Spectrometry Interactive Virtual Environment (MassIVE) database (Identifier: MSV000081382) [3].

## 6 Acknowledgements

This work is supported by grants from National Key Research and Development Project Subject (Grant No. 2021YFA1301603), “Pioneer” and “Leading Goose” R&D Program of Zhejiang (2024SSYS0035), the Zhejiang Province Leading Geese Plan (2024SSYS0035), and the Key Research and Development Program of Zhejiang Province (Grant No. 2022C03037). We gratefully acknowledge the support of Westlake University HighPerformance Computer Center and Shanghai Artificial Intelligence Laboratory. We thank the Research Center for Industries of the Future (RCIF) at Westlake University for partially supporting this work. We thank Dr. Yingying Sun for generously providing the raw files of several benchmark datasets, and our laboratory colleagues for their support and assistance during the data download process.

## 7 Author contributions

T.G. and S.S. co-directed and supported the study. J.A., Xiaofan Z., and T.Z. established the MassNet dataset. Xiaofan Z. developed the tensor format (MSDT) for storing database search results and spectral data. J.A. prepared the 15-species benchmark dataset. Xiang Z. served as the principal developer of XuanjiNovo, with algorithm design guidance from S.S. Xiang Z. and J.W. carried out *de novo* sequencing training and inference across all datasets. Z.G., J.W., Y.D., P.L., and Z.N. contributed to interpretive analyses. The manuscript was co-written by J.A., Xiang Z., and J.W., with all authors involved in reviewing, revising, and approving the final version.

## 8 Competing interests

T.G. is a shareholder of Westlake Omics Inc. The remaining authors declare no relevant competing interests.

## 9 Supplementary information

**Table S1**. Summary of PSMs, peptide precursors, and identified proteins by species.

**Table S2**. Detailed information of the 15-species benchmark dataset.

**Supplementary Figure 1.**
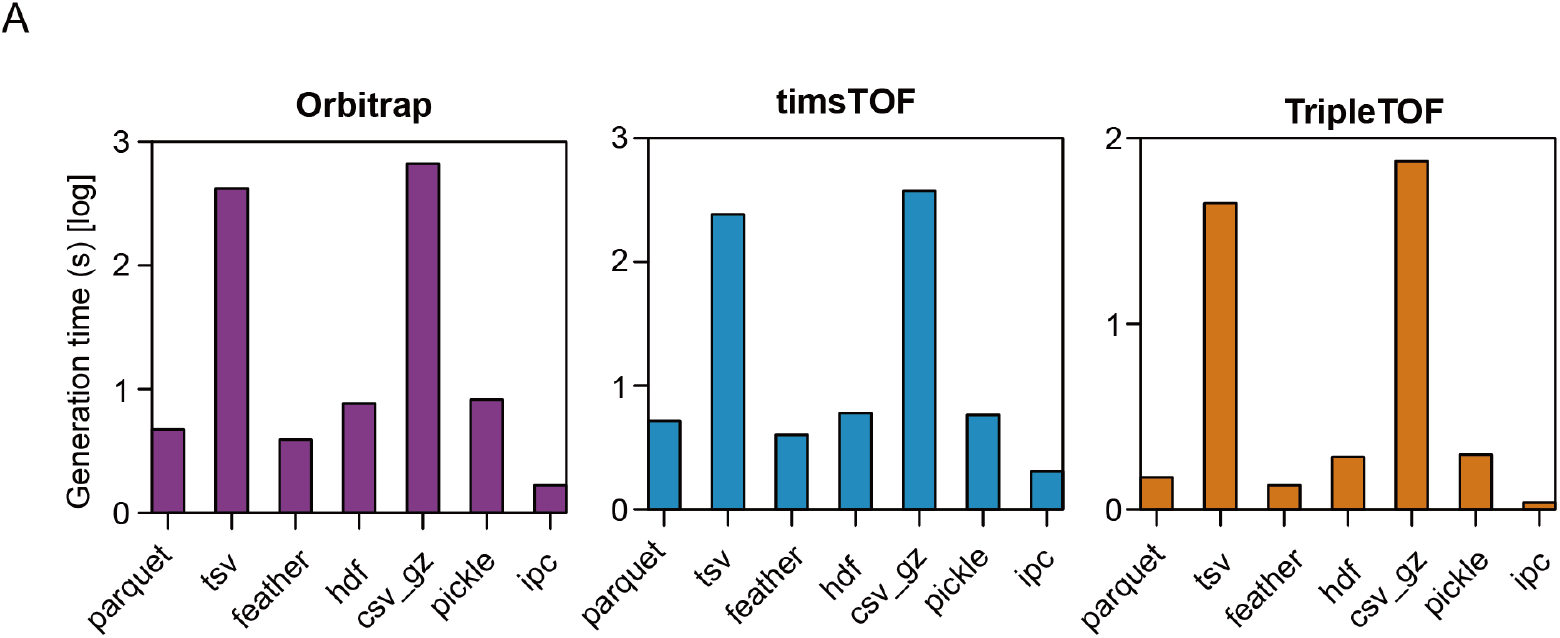
Generation time of various serialization formats across different instrument platforms. (A) Log-scaled generation time (in seconds) for seven data serialization formats including parquet, tsv, feather, hdf5 (hdf), csv gz (gzipped), pickle, and ipc, evaluated on three mass spectrometry instrument types: Orbitrap (purple), timsTOF (blue), and TripleTOF (orange). For each type of mass spectrometer, at least six raw files were randomly selected, and the average generation time for each format was calculated.

**Supplementary Figure 2.**
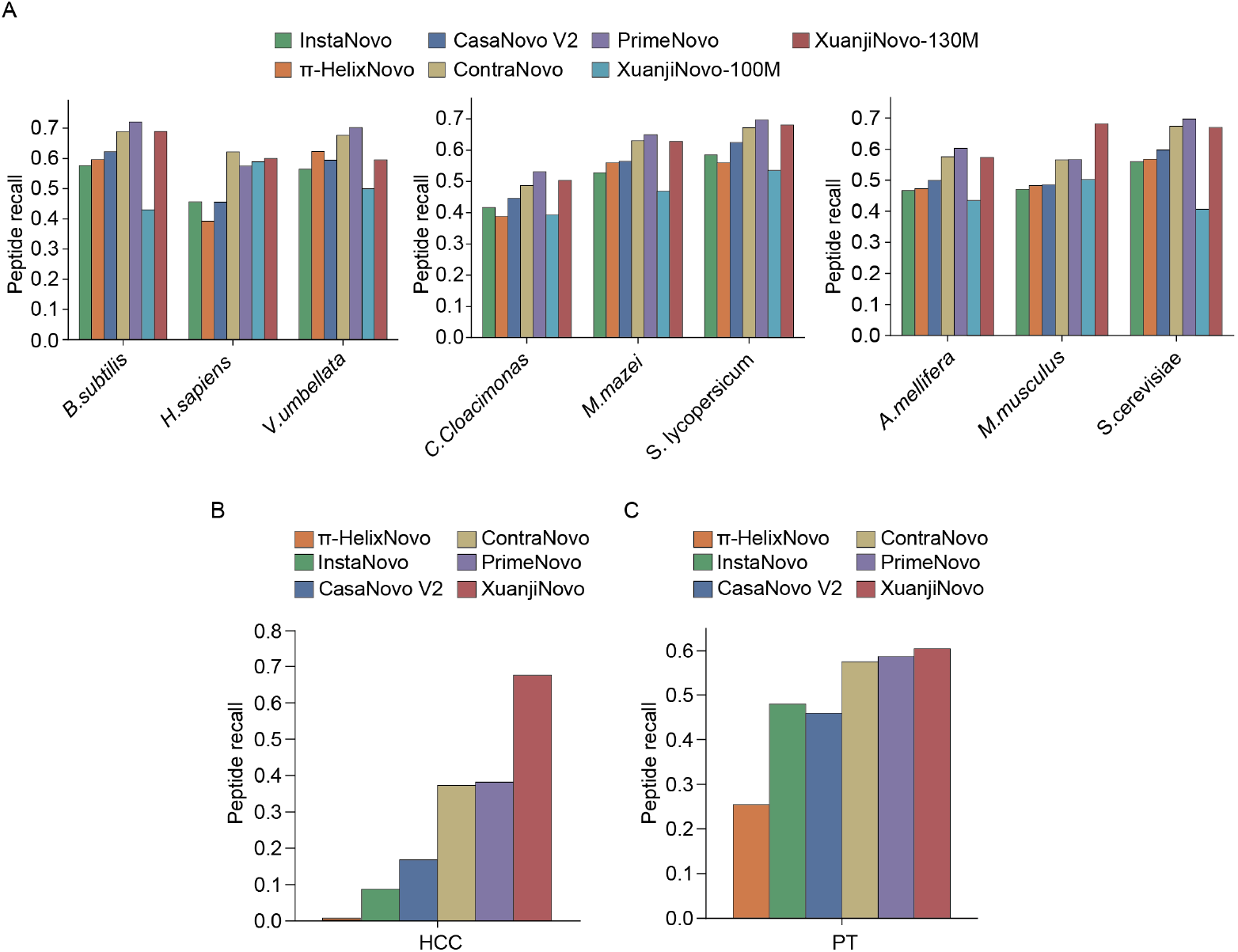
Evaluation of *de novo* peptide sequencing performance on other benchmark datasets. (A) Peptide recall across nine species in the 9-species benchmark dataset using seven models: InstaNovo, *π*-HelixNovo, ContraNovo, CasaNovo V2, PrimeNovo, XuanjiNovo-100M, and XuanjiNovo-130M. (B) Peptide recall on the HCC dataset. (C) Peptide recall on the PT dataset. The *π*-HelixNovo model is publicly available and was trained on datasets derived from eight species, excluding human.

## Notes

### Competing Interest Statement

The authors have declared no competing interest.

